# Transcriptional signatures of connectomic subregions of the human striatum

**DOI:** 10.1101/089458

**Authors:** Linden Parkes, Ben Fulcher, Murat Yücel, Alex Fornito

**Author notes:** Corresponding author, 770 Blackburn Road, Clayton, Victoria 3168, Australia. Abbreviated title: Connectivity and gene expression in striatum. Author contributions: L. P., B. F., and A. F. designed research; L. P. performed research; L. P., B. F., and A. F. contributed unpublished reagents/analytic tools; L. P., B. F., and A. F. analyzed data; L. P., B. F., M. Y., and A. F. wrote the paper.

## Abstract

Functionally distinct regions of the brain are thought to possess a characteristic connectional fingerprint – a profile of incoming and outgoing connections that defines the function of that area. This observation has motivated efforts to subdivide cortical and subcortical areas using their patterns of connectivity. However, it remains unclear whether these connectomically-defined subregions of the brain can be distinguished at the molecular level. Here, we combine high-resolution diffusion-weighted magnetic resonance imaging with comprehensive transcriptomic data to show that connectomically-defined subregions of the striatum carry distinct transcriptional signatures. Using data-driven clustering of diffusion tractography, seeded from the striatum, in 100 healthy individuals, we identify a tripartite organization of the caudate and putamen that comprises ventral, dorsal, and caudal subregions. We then use microarray data of gene expression levels in 19 343 genes, taken from 98 tissue samples distributed throughout the striatum, to accurately discriminate the three connectomically-defined subregions with 80-90% classification accuracy using linear support vector machines. This classification accuracy was robust at the group and individual level. Genes contributing strongly to the classification were enriched for gene ontology categories including dopamine signaling, glutamate secretion, response to amphetamine, and metabolic pathways, and were implicated in risk for disorders such as schizophrenia, autism, and Parkinson’s disease. Our findings highlight a close link between regional variations in transcriptional activity and interregional connectivity in the brain, and suggest that there may be a strong genomic signature of connectomically-defined subregions of the brain.

## 1 Introduction

The functional specialization of specific brain regions is governed, in part, by the anatomical connectivity of that area with the rest of the brain (Passingham et al., 2002). The idea that distinct functional areas of the brain may possess a unique “connectional fingerprint” has motivated a large body of work that attempts to parcellate the brain into functional zones based on regional variations in connection profiles (Behrens and Johansen-Berg, 2005, Eickhoff et al., 2015). Connectivity-based parcellation has been used to examine the subregional architecture of cortical areas such as the medial frontal cortex (Johansen-Berg et al., 2004), lateral premotor cortex (Tomassini et al., 2007), and Broca’s area (Anwander et al., 2006), as well as subcortical areas such as the subthalamic nucleus (Lambert et al., 2012), insula (Alcauter et al., 2015), thalamus (Behrens et al., 2003), and striatum (Bohanna et al., 2011), as well as the entire brain (Fan et al., 2016). While these connectivity-based parcellations have been successful in defining functionally relevant subregions of the brain (e.g., Fan et al., 2016, Gordon et al., 2016), it remains unclear whether connectomically-defined subregions show variation at the molecular level. Addressing this question is critical for understanding the relationship between micro-scale and macro-scale brain organization.

The recent availability of large-scale transcriptomic data, incorporating measures of the expression of thousands of genes measured throughout the brain (Shen et al., 2012, Hawrylycz et al., 2012, Oldham et al., 2008), has afforded an unprecedented opportunity to investigate the relationship between gene expression and brain connectivity. Preliminary work has shown that regional variation in gene expression across the brain can be used to delineate major divisions of the cortex, such as the visual, motor, frontal, and sensory cortices, in both the mouse (Hawrylycz et al., 2010) and human brains (Hawrylycz et al., 2012, Oldham et al., 2008). Furthermore, a close link has been demonstrated between transcription levels and inter-regional structural and functional connectivity (French and Pavlidis, 2011, Fulcher and Fornito, 2016, Kaufman et al., 2006, Krienen et al., 2016, Wolf et al., 2011), suggesting that regional variations in extrinsic connectivity are closely tied to gene expression patterning.

The striatum has been a popular target for connectivity-based parcellation (Draganski et al., 2008, Bohanna et al., 2011, Lehéricy et al., 2004, Leh et al., 2007, Tziortzi et al., 2014). A large body of evidence indicates that the striatum can be delineated into distinct functional zones along a rostroventral to dorsocaudal gradient. According to one prominent, tripartite model of striatal organization (Parent and Hazrati, 1993, 1995, Haber, 2003), ventral areas of the striatum play an important role in emotional function and are preferentially connected to limbic areas such as the orbitofrontal cortex, amygdala and hippocampus; dorsal areas support higher-order associative processing and connect predominantly with dorsolateral and prefrontal regions; and caudal areas mediate sensorimotor processes, displaying strong connectivity with motor and sensory cortices. These subdivisions have been differentially implicated in different disorders, including obsessive-compulsive disorder (Harrison et al., 2009), addiction (Everitt and Robbins, 2013), Parkinson’s disease (DeLong and Wichmann, 2007), psychosis (Fornito et al., 2013, Dandash et al., 2014), and problem gambling (Koehler et al., 2013). These subdivisions also differ in terms of their chemoarchitecture. In situ hybridization of human tissue samples (Gurevich, 1999), immunochemistry in rats (Levey et al., 1993), and microarray analysis in mice (Olsen et al., 2008) all suggest that dopamine receptors are differentially expressed across different striatal subregions. For example, while D2 receptors are more strongly expressed in the caudate and putamen compared to the nucleus accumbens (Levey et al., 1993, Olsen et al., 2008), the opposite pattern is observed for D3 receptors (Gurevich, 1999). Furthermore, D2 receptor concentrations within the caudate and putamen decline along a rostrocaudal gradient, being lower in caudal regions (Levey et al., 1993). These findings indicate that the density of distinct classes of dopamine receptors varies across different subregions of the striatum. However, previous work has had to trade off spatial coverage of the striatum against complete coverage of the genome. In other words, investigators typically examine the expression of specific subsets of genes across large swathes of the striatum (e.g., Levey et al., 1993, Gurevich, 1999), or measure expression levels in tens of thousands of genes at only a few spatial locations (e.g., Olsen et al., 2008).

The recently published Allen Human Brain Atlas (AHBA; Hawrylycz et al., 2012), which uses microarray probes to assay expression levels across almost the entire genome, measured in 3702 tissue samples distributed throughout the brain, opens new opportunities to comprehensively characterize molecular function in the striatum. In this study, we used these data to characterize the transcriptional signature of connectomically-defined subregions of the human striatum. We first reconstructed the connectivity of each of over 2000 voxels in the left and right striatum with 86 cortical and subcortical brain regions covering the whole brain, and used data-driven clustering to identify connectomically-defined striatal subregions. We then used linear support vector machines to determine whether we could accurately classify which connectomic subregion each of 98 striatal tissue samples contained in the AHBA was drawn from, based on the transcriptional profile of that sample. This approach allowed us to comprehensively evaluate transcriptional distinctions between different striatal subregions. In particular, it allowed us to evaluate whether variations in dopaminergic function are indeed a major molecular correlate of striatal organization, or whether other neurotransmitter or physiological systems also play a role.

## 2 Materials and Methods

We combined high-quality DWI data from the HCP with multivariate clustering techniques to produce an objective, data-driven parcellation of the striatum. We then used high throughput gene expression data from the AHBA to investigate whether connectomically-defined striatal subregions display distinct patterns of gene expression. Figure 1 provides an overview of the steps involved in these analyses.

**Figure 1.**
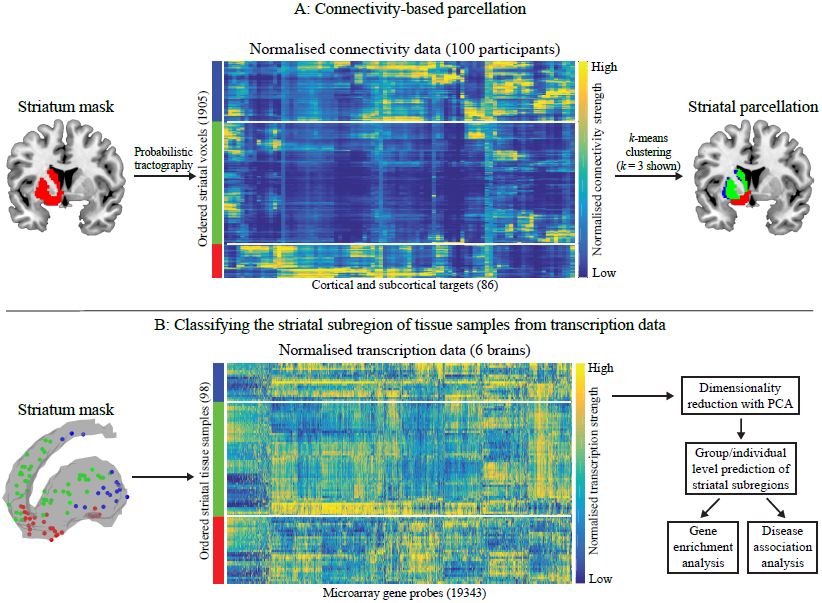
A schematic illustration of the two key analyses of the study. (A) An outline of the steps involved in connectomic parcellation of the striatum. Probabilistic tractography was performed from each striatal voxel to 86 cortical and subcortical targets for 100 participants. Connectivity data was averaged across the 100 participants and normalised. Striatal voxels were then grouped together based on similar connectivity profiles using k-means clustering. Results of *k*-means clustering are represented by the colourbar on the left. Blue corresponds to voxels in cluster 1, green to cluster 2, and red to cluster 3. This colour scheme is carried forward to brain space on the right. For visualisation, targets (columns) have been reordered by hierarchical clustering to place targets with similar connectivity to striatal voxels close together, and voxels (rows) have been ordered according to the *k*-means (*k* = 3 shown here) clustering (left colourbar) as well as reordered within clusters according to a hierarchical clustering to place voxels with similar connectivity close to one another. (B) An outline of the steps involved in genomic cross-validation of the connectomic parcellation using transcription data from the AHBA. A total of 98 tissue samples within the striatum were extracted from the 6 brains in the AHBA. Tissue samples are colour coded according to which connectomically-defined striatal subregion they belong to (left colourbar and circles in striatum mask). Transcription data for 19 343 gene probes were retained. For visualisation, transcription data was normalised before being reorganised according to the same steps outlined above. Principal component analysis (PCA) reduced the dimensionality of the microarray data before performing support vector machine classification (SVM) analyses at the group and individual levels. Finally, gene enrichment and disease association analyses were performed on select PCs shown to discriminate striatal subregions.

### 2.1 Data

DWI data from 100 unrelated healthy participants (54 males, 46 females, age range of 22-35 years) were downloaded from the HCP database (Van Essen et al., 2013). We used the minimally preprocessed DWI and structural data, the full details of which can be found elsewhere (Glasser et al., 2013). In brief, the HCP data were acquired using a customized Siemens 3T Skyra MRI machine. The DWI data (1.25 mm isotropic voxels, TR = 5520 ms, TE = 89.5 ms, flip angle = 78 degrees, echo spacing = 0.78 ms) were obtained using a multi-shell protocol that allowed for concurrent acquisition of interspersed diffusion weighted gradients (*b* = 1 000, 2000, and 3000 s/mm^2^). Acquisition of reverse phase encoding b = 0 pairs allowed for the estimation of the underlying magnetic field inhomogeneities that caused geometric distortions in the EPI images, and their subsequent correction (Andersson et al., 2003). Additional field inhomogeneities caused by head motion were also corrected during this step. T1-weighted structural data were additionally collected for each participant (0.7 mm isotropic voxels, TR = 2400 ms, TE = 2.14 ms, flip angle = 8 degrees). Gradient nonlinearities in the DWI data produced by the customized head coil used in the HCP were corrected before the DWI data were registered to native T1 volume space. Diffusion gradients were rotated accordingly. Diffusion tractography (see below) was performed in each participant’s T1 space.

### 2.2 Generation of seed and target areas for tractography

Initial attempts to parcellate the striatum based on diffusion data have examined structural connectivity in relation to cortical lobes, which may mask finer-grained distinctions between different striatal areas (Draganski et al., 2008). These studies also defined subregion boundaries by selecting the connection with the highest probability for each voxel (a winner-take-all approach), regardless of the relative strength of other connections (Lehéricy et al., 2004, Bohanna et al., 2011, Tziortzi et al., 2014). Given the detailed connectivity profiles apparent within striatal voxels (Fig. 1A), focusing only on the maximal probability connection may overlook the potential for striatal subregions to have complex and diverse connectivity profiles. Our aim was to incorporate these multifaceted connectivity profiles into the parcellation analysis.

We took the caudate, putamen, and nucleus accumbens, as defined in the Desikan-Killiany atlas (Desikan et al., 2006) provided by the HCP, in each hemisphere and combined them to form a pair of seeds defined at 2mm3 voxel resolution for tractography (i.e., a left and a right hemisphere striatum mask). There were several targets used for this study: (i) 68 cortical areas (34 per hemisphere); (ii) left and right hippocampus and amygdala; and finally (iii) seven subregions of the thalamus defined according to cortico-thalamic structural connectivity (Behrens et al., 2003). Target sets (i) and (ii) were both defined according to the Desikan-Killiany parcellation. The segmented thalamus mask was chosen because different striatal subregions connect topographically to distinct thalamic nuclei (Giménez-Amaya et al., 1995).

### 2.3 Probabilistic tractography

Analysis of the DWI data was performed using the Functional Magnetic Resonance Imaging of the Brain (FMRIB) Software Library (FSL). Specifically, tractography was performed using FMRIB’s diffusion toolbox (FDT) (http://fsl.fmrib.ox.ac.uk/fsl/fslwiki/) (Behrens et al., 2003). The fibre orientation density function was modelled for the DWI data on a voxel-wise basis using a ball & stick model developed for use with multi-shell datasets (Jbabdi et al., 2012). Streamline tracking was performed from each voxel in a given seed mask using a probabilistic algorithm. The advantage of utilizing multi-shell acquisitions is that they increase the sensitivity of detecting crossing streamlines (Sotiropoulos et al., 2013). Also, FSL’s tractography method can simultaneously handle surfaces and volumes as regions of interests, which allows for more accurate determination of streamline termination points compared to just using volumes.

For each participant, FDT’s bedpostx function was used to estimate streamline orientation and uncertainty at each voxel in the DWI data by estimating a distribution of diffusion parameters using Markov Chain Monte Carlo sampling, with 3 fibers modelled per voxel, Rician noise assumed (Sotiropoulos et al., 2013), and taking into account the gradient nonlinearities generated by the customized scanner used by the HCP. FDT’s probtrackx2 calculated the probability of connectivity between each seed voxel with each target using 5000 streamlines per voxel, a 0.2 curvature threshold, and loopcheck termination (a step designed to terminate streamlines that loop back on themselves). Included with the HCP data are transformation matrices which provide a spatial mapping from a participant’s T1-weighted/diffusion space to MNI space. These matrices enabled us to perform tractography in each participant’s diffusion space with ROIs defined in MNI space and then store the results in MNI space. Tractography was run from each of the 1923 seed voxels in the left striatum and the 1905 seed voxels in the right striatum to the 86 bilateral cortical and subcortical targets outlined above. The results were represented as separate 1923 × 86 and 1905 × 86 connectivity matrices, where each element was the count of streamlines that intersected a given seed voxel and target. Streamline counts for each voxel-target pair were averaged across subjects separately for the left and right hemispheres.

### 2.4 Connectivity-based parcellation of the striatum

We parcellated the striatum masks by grouping seed voxels with similar profiles of extrinsic whole-brain connectivity (Fig 1A) using *k*-means clustering (Hastie et al., 2009). Each seed voxel was represented in terms of its connectivity profile as an 86-feature vector, where each feature was a count of the number of reconstructed streamlines that intersected each of the 86 tractography targets. Note that we use the term “target” here to refer to a non-striatal region used as a target for tractography - not to imply that the region was the target of an efferent projection from the striatum (since afferent and efferent projections cannot be resolved with diffusion MRI). ROI volume varied across the 86 connectivity targets used for our tractography and larger targets are more likely to intersect streamlines than smaller targets. To correct for this target volume bias, streamline counts were normalized for each connectivity target using the sigmoidal function, *S*(*x*) = [1 + exp(−(*x* − *μ*)/*σ*)]−^1^ where *μ* is the mean and *σ* is the standard deviation of streamline counts across all seed voxels for a given target. Values were then scaled linearly to the unit interval to yield values between 0 (low relative connectivity to the target) and 1 (high relative connectivity to the target). Alternative normalization methods, such as *z*-score, yielded qualitatively similar results. Using the kmeans function in Matlab 2014b, the dissimilarity between the normalized connectivity profiles for a given striatal voxel was quantified using the squared Euclidean distance metric. In turn, distances are used in the *k*-means clustering algorithm to form groups of similar connectivity profiles.

Given that our primary aim was to examine the gene transcriptional signatures of striatal subregions, and given the limited anatomical coverage of tissue samples in the AHBA (see sections 2.5 and 2.5.1 below), we began our analysis by partitioning the striatal seed voxels into three homogeneous clusters (*k* = 3). However, due to evidence for finer connectivity-based parcellations of the striatum (e.g., Draganski et al., 2008, Baliki et al., 2013, Piray et al., 2015) we also investigated higher levels of *k* (*k* = 4, 5, 6) to examine the rostrocaudal gradient of striatal organization in more detail. For a given *k*, the *k*-means clustering algorithm yields a labelling of each seed voxel in the left and right striatum masks that represents that voxel’s allocation to one of the *k* clusters.

#### 2.4.1 Individual differences in striatal organization

Our group-level delineation of striatal subregions was based on connectivity data averaged across a sample of 100 individuals. To examine whether this group-level representation is an accurate summary of striatal organization at the level of individuals, we reproduced the *k* = 3 clustering solution for each participant separately (using the same method outlined in Section 2.4) and quantified the similarity between these individual-level parcellations and the group-level parcellation. To ensure our results were not biased, we also quantified the similarity between the individual-level parcellations and group-level parcellations generated using the *N*-1 remaining participants. Finally, to assess inter-individual variability in striatal demarcations, we performed pairwise comparisons between the individual-level parcellations. The similarity between parcellations was quantified using the normalized variation of information (VI) (Meilă, 2007, Rubinov & Sporns, 2011), which is an information theoretic measure of the overlap between two clustering solutions. Values of the normalized VI range between zero and one, where zero corresponds to identical clustering solutions and one corresponds to maximally different clustering solutions.

#### 2.4.2 Extrinsic connectivity of striatal subregions

Voxels within each striatal subdivision identified using *k*-means clustering should exhibit a homogenous profile of extrinsic connectivity with the rest of the brain that is distinct from the other subdivisions. In order to characterize the distinctive pattern of extrinsic connectivity for each identified striatal cluster, we quantified the relative strength of connectivity between each striatal cluster in the *k* = 3 solution and the connectivity targets using *t*-tests. We first calculated *z*-scores of the connectivity data described in section 2.4.1 across all the individual participants to ensure mean connectivity of zero for each target, which allowed us to conduct single sample *t*-tests on the connectivity between each cluster and each target. Second, to ensure identical clustering across participants, we applied the *k* = 3 solution generated at the group level to each participant’s connectivity data and averaged across voxels within each cluster for each target. This yielded a 3 × 86 (cluster × target) connectivity matrix for each participant representing the mean connectivity between each cluster and each target for a given hemisphere. Third, we performed one-sample *t*-tests for each cluster to identify the connections that had mean connectivity significantly greater than zero across all participants. This resulted in significant cluster to target connections for each striatal subdivision in both hemispheres. For visualization, we summed the significant *t*-values for a given target from each hemisphere to generate a single *t*-value for each cluster and target. This analysis resulted in three maps that revealed the predominant connectivity targets for each striatal cluster, collapsed across hemispheres.

### 2.5 Gene expression

To investigate whether the connectomically-defined striatal subregions were associated with distinct transcriptional profiles, we analyzed microarray data from the AHBA, the full details of which can be found elsewhere (Hawrylycz et al., 2012). In brief, the AHBA includes normalized genome-wide microarray data (58 692 probes) for 3702 tissue samples taken from the cortex and subcortex of six post-mortem adult brains (approximately 400-500 tissue samples per brain). Each has an associated stereotaxic coordinate in MNI space. We focused our genetic analysis on the left hemisphere because four of the six brains in the AHBA database contained no samples in the right hemisphere, and the two remaining brains did not contain a sufficient number of samples to allow for a subregional analysis of the right hemisphere.

There was a large amount of redundancy in the microarray data provided by the AHBA. To ensure each microarray probe assayed a unique gene, probes that did not carry an entrez identification number were removed, and in cases where certain genes were assayed by multiple probes, a single probe was selected by performing principal component analysis (PCA) on the duplicate probes and retaining the probe with the highest loading on the first principal component (PC). Since we are interested in the transcriptomic profiles of the striatum, we next filtered the 3702 tissue samples by matching the MNI coordinates supplied by the AHBA with the left hemisphere striatum mask from the connectivity analysis. This was done in two steps. First, we converted our binary striatal mask (see section 2.2) to a 3-D shape and retained only those tissue samples that were located within this shape using Matlab’s alphaShape and inShape commands, respectively. Second, we found the shortest Euclidean distance between the MNI coordinates of each striatal tissue samples and the MNI coordinates of each of the voxels in our parcellation. Thus, a final 98 × 19 343 tissue sample × gene expression probe data matrix was obtained.

#### 2.5.1 Genomic classification of striatal subdivisions

We used the AHBA data to examine whether transcription data could be used to accurately classify 98 striatal tissue samples into one of three connectomically-defined striatal clusters. The 98 tissue samples were matched to a striatal cluster label based on the MNI coordinates provided by the AHBA. We then used a support vector machine classifier (SVM) with a linear kernel to classify the tissue samples for each of the striatal clusters (Hastie et al., 2009). SVM classifiers can only resolve classification problems with more than two groups by performing multiple separate two-class classifiers. Our primary goal here is to determine whether gene expression can distinguish any given striatal subregion from the others, so each SVM was specified as a one vs. all others problem (Hastie et al., 2009). In other words, we tested whether cluster 1 could be distinguished from clusters 2 and 3, whether cluster 2 could be distinguished from clusters 1 and 3, and whether cluster 3 could be distinguished from clusters 1 and 2. This procedure was repeated for each of the three clusters. Ten-fold cross-validation with 50 repeats was used to obtain out-of-sample estimates of classification performance, measured as the overall classification accuracy (proportion of correctly classified tissue samples). To ensure robust estimates of classification accuracy, new random stratified subsets of tissue samples were used as testing data for each repetition.

Using the full AHBA dataset in the SVM classifier would result in 19 343 features (gene expression measures), with only 98 observations (striatal tissue samples). In order to minimize the potential for overfitting the data (Hastie et al., 2009) and find an interpretable representation of the transcription data, we reduced this feature space using PCA. In contrast to mass univariate testing of subregional gene expression differences, using PCA allows us to characterize the multivariate genomic profiles for each striatal subregion, providing an estimate of multivariate discriminability. We retained the 10 leading PCs, which explained 67% of the variance in the data (additional PCs each accounted for <2% of the variance in the data and did not yield classification accuracies higher than an empirical null distribution). We ran the SVM separately for each PC, and separately tested for an improvement in classification accuracy by cumulatively adding each PC until all ten were used as features in the classifier. The statistical significance of the obtained classification accuracies was quantified using permutation testing by repeating the above classification procedures 5 000 times, each time independently shuffling the striatal subregion allocation of the tissue samples to generate an empirical null distribution for each cluster separately. Calculating separate null distributions is necessary because SVM classification was performed on separate striatal clusters, each with a differing number of tissue samples assaying transcription levels.

Due to limited coverage of the striatum in the AHBA, genomic classification of striatal clusters was only possible for the *k* = 3 solution obtained from the *k*-means clustering of the connectivity data. Moving to *k* = 4 would result in only 11 samples being included in some regions, which was not sufficient for robust genomic classification. Similarly, our genomic analyses were restricted to the left hemisphere due to insufficient coverage of the right striatum in the AHBA (e.g., for the *k* = 3 solution in the right hemisphere, the smallest cluster contained only a single sample).

As a result of uneven tissue sample coverage across the striatum and variation in the size of each striatal cluster, the AHBA tissue samples were distributed unevenly across the clusters. To ensure that the SVM classifier was not biased by this class imbalance, we weighted the misclassification cost of each of the tissue samples according to the inverse probability of being located within that striatal cluster. Specifically, we assigned all tissue samples belonging to the striatal cluster, i, a weight, *W*_*i*_ = 1/*p_i_*, where *p_i_* is the proportion of tissue samples in cluster *i* (Liu et al., 2008).

#### 2.5.2 Gene ontology enrichment analysis

The PCA of the microarray data yields a set of PC coefficients that measure the weighting of each gene’s expression levels onto a given PC. To see if gene probes with a higher loading onto certain PCs were associated with particular biological processes, we performed an enrichment analysis of GO categories (Ashburner et al., 2000) using Gene Score Resampling (GSR) implemented in the ErmineJ software package (Lee et al., 2005). We performed GSR separately for each PC, using the absolute PC coefficients as gene scores, yielding a *p*-value for each GO category. We used 10^7^ iterations with full resampling for each iteration, considered gene set sizes between the range of 5-100, and took the mean scores in a GO group as a summary statistic. Significance levels were adjusted for multiple comparison testing using the Benjamini-Hochberg False Discovery Rate (FDR) correction (Benjamini and Hochberg, 1995). GSR has the potential to yield long lists of significant GO categories. In order to reduce the number of significant GO categories and facilitate interpretation, where necessary, REVIGO was used to remove redundant GO terms from the results (Supek et al., 2011). REVIGO works by selecting a representative GO term from related categories in the GO hierarchy (e.g., by parent, child, or sibling connections) using the SimRel algorithm (Schlicker et al., 2006).

#### 2.5.3 Inter-individual variability of AHBA tissue samples

To maximize the number of tissue samples spanning the striatal subdivisions, we pooled transcription data across all six brains in the AHBA. However, since these six subjects are from different age groups, genders, and ethnicities, we also conducted a second analysis designed to investigate the effect of this individual variability on the results obtained using the pooled data. Specifically, we performed individual-level classification analyses using a “leave one brain out” approach. Instead of using a random stratified subset of tissue samples, we trained the SVM classifiers on tissue samples from five of the six brains and tested on tissue samples from the remaining brain. We repeated this process until each brain had been left out and classified. To prevent the testing data influencing the training data in any way, the PCA was run for each iteration on the training data and the test set was transformed into this PC space.

#### 2.5.4 Disease association analysis

In addition to looking for biological processes enriched in transcriptional PCs, we also explored whether the PCs were associated with particular diseases using the Autworks database (http://autworks.stanford.edu/) (Nelson et al., 2012). Autworks comprises evidence-based disease-gene annotations across the whole human genome for thousands of diseases. Hence, each disease has an associated gene set. Autworks accepts user-defined gene sets and computes the probability that a user-defined gene set overlaps with each of the inbuilt disease gene sets. We took sets of genes that had the strongest correlations (top 10% as in Hawrylycz et al., 2015) with each PC to determine whether any of our PCs were over-represented in gene sets involved in disease. Taking the top 10% of genes keeps the size of the gene sets constant across PCs. This is important because PCs with longer gene sets can show enrichment of more categories, simply because a larger number of genes has been analyzed. Nonetheless, to evaluate the robustness of our results, we cross-validated our findings by selecting genes that correlated significantly with each PC using a permutation test. With this approach, genes that were significant at *p*<0.05, (permutation test, FDR corrected) were retained as gene sets. This method is a statistically principled framework for selecting PC-related genes, but can vary in the number of genes included across gene sets. Thus, larger gene sets are more likely to be implicated in a broader range of diseases. Comparing results from the two methods allows identification of the most robust associations with disease.

#### 2.5.5 Validation against other striatal parcellations

A connectivity-based parcellation based on *k*-means clustering is but one of many potential methods for defining striatal subregions. To determine whether other approaches to parcellation yield similar transcriptional signatures, we repeated our analysis using two alternative, publicly available striatal atlases. The first delineates the caudate, putamen and ventral striatum (including nucleus accumbens) based on anatomical boundaries, as defined in the Desikan-Killiany atlas (Desikan et al., 2006. see section 2.2). The second was derived using an alternative connectivity-based parcellation strategy, in which striatal voxels were clustered based on the highest connection probability (winner-takes-all approach) to a small set of cortical targets (defined at the lobar level) rather than on multivariate connectivity profiles (Tziortzi et al., 2014). The former atlas is freely used in the Freesurfer software package and the latter is provided with FSL.

### 2.6 Results

#### 2.6.1 Connectomic parcellation of the striatum

Our primary aim was to use connectional fingerprints to identify striatal subregions in a data-driven way, and then to examine how subregional functional differences are expressed at the molecular level. We clustered individual striatal voxels according to their patterns of extrinsic connectivity with the rest of the brain. When extracting three clusters (i.e., *k* = 3), the results identify clear, anatomically contiguous subregions that encompass the ventral, dorsal, and caudal portions of the striatum in both hemispheres (Fig. 2A). The caudal cluster (blue in Fig. 2A) encompasses the lateral putamen, extending caudally, and the tail of the caudate. The dorsal cluster (green in Fig. 2A) contains most of the head and dorsomedial sections of the caudate and the medial putamen. Finally, the ventral cluster (red in Fig. 2A) consists primarily of the nucleus accumbens as well as ventral portions of the putamen. For the remainder of this paper we refer to the three striatal clusters using this ‘caudal’, ‘dorsal’, ‘ventral’ naming scheme.

**Figure 2.**
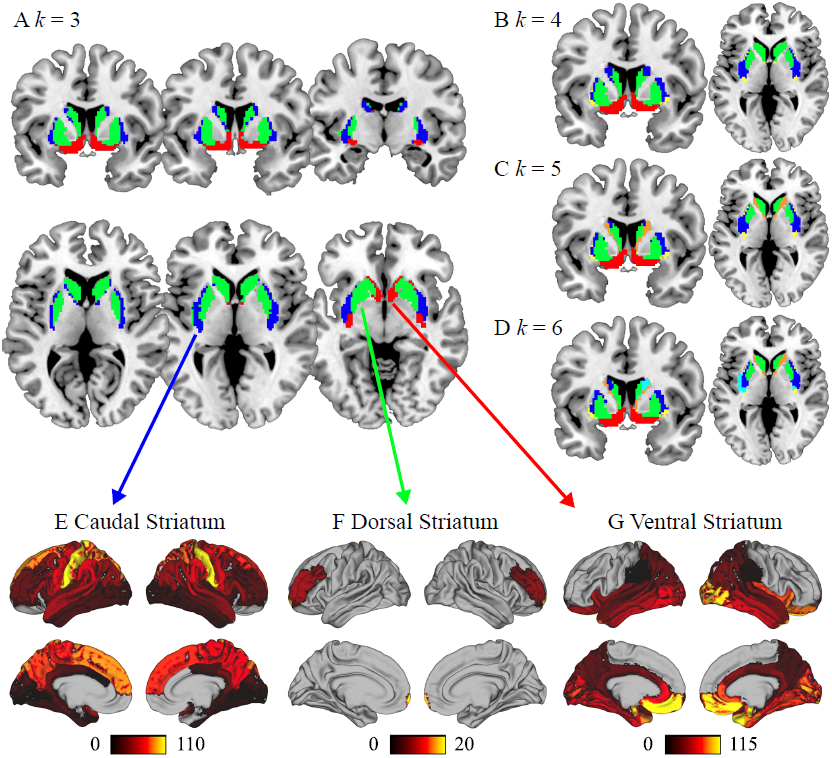
Parcellation of the human striatum based on whole-brain structural connectivity. Subregions of the tripartite striatum show distinct cortical connectivity profiles that are consistent with their functional roles. (A) Using *k*-means clustering, with *k* = 3, to demarcate the striatum based on connectivity with 86 cortical and subcortical targets covering the whole brain, yields a ‘caudal’ cluster (blue), a ‘dorsal’ cluster (green), and a ‘ventral’ cluster (red). (B-D) At higher levels of *k*, striatal clusters subdivide along mediolateral and rostrocaudal gradients. (E-G) Connectivity data for all participants were *z*-scored and averaged across voxels within each cluster for each target in each hemisphere separately. Single sample/-tests were performed to determine significant cluster to target connectivity for each hemisphere. Significant absolute/-values (p<0.05 FDR corrected) were averaged across the left and right hemispheres and plotted on the surface of the brain to visualise the specific patterns of connectivity characteristic of each striatal cluster. (E) The caudal striatum shows strong connectivity with sensorimotor areas of the cortex. (F) The dorsal striatum shows connectivity to frontal regions. (G) The ventral striatum shows strong connectivity to limbic areas.

We also parcellated the striatum at higher levels of *k* to examine finer distinctions in the rostrocaudal gradient. The results are shown in Figure 2B. For each additional level of *k*, the caudal, dorsal and ventral subregions subdivide along a mediolateral and rostrocaudal gradient, consistent with prior findings (Reep and Corwin, 1999, Draganski et al., 2008). Note that there is no sudden change in boundary locations, consistent with hierarchical anatomical organization.

#### 2.6.2 Individual differences in striatal organization

The data-driven grouping of striatal voxels was generated at the group level (Fig. 2A) and may not be representative of striatal organization at the individual level. Thus, we generated a *k* = 3 solution for each participant separately and quantified the pairwise discrepancy between participants, as well as the discrepancy between each participant and two group-level parcellations (i.e., generated with and without the participant’s own connectivity data) using the variation of information (VI, see methods). Comparing each participant’s *k* = 3 solution to the group-level *k* = 3 solution revealed an average VI of 0.11 ± 0.02 (mean ± SD) for both the left hemisphere and the right hemisphere regardless of whether the group-level solution was generated including the subject’s connectivity data or not. Additionally, comparing participants' *k* = 3 solutions in a pairwise manner revealed an average VI of 0.13 ± 0.02 for both the left and the right hemispheres. The low mean VI and small standard deviation indicates that there is a large amount of overlap between the *k* = 3 solution generated at the group level and *k* = 3 solutions generated at the individual level, and that the parcellation is quite robust across the individual participant solutions. In turn, this suggests that the group level solution was a good representation of striatal organization at the individual level for both hemispheres.

#### 2.6.3 Extrinsic connectivity of the caudal, dorsal and ventral striatal subregions

Having identified three distinct striatal subdivisions on the basis of extrinsic connectivity, we wanted to understand the specific connectivity profile of each cluster in the *k* = 3 solution. We mapped the anatomical distribution of the *t*-values indexing group connectivity with each target (see section 2.4.2). Targets characteristic of each striatal cluster (*p*<0.05, FDR corrected), are shown in Figure 2E–G. Note that these figures represent statistical maps generated for each cluster separately and do not illustrate connectivity targets that are more connected to one cluster compared to another. Furthermore, the absence of a statistically significant connection between cluster and target within these maps indicates that the connectivity for all participants, represented as *z*-scores and averaged across voxels within each cluster for each target, was not significantly different from zero in this sample; not that an anatomical connection between these areas was absent.

The caudal striatal cluster (Fig. 2E) had widespread significant connectivity with 70 and 67 targets in the left and right hemisphere, respectively. Connectivity was strongest with the sensory and motor cortices, as well as superior frontal and parietal regions. The caudal cluster also showed significant connectivity to the thalamus and hippocampus. By contrast, the dorsal striatal cluster (Fig. 2F) showed a more focal profile of connectivity, with significant projections to the middle frontal gyri and the frontal poles. Finally, the ventral striatal cluster (Fig. 2G) showed an expansive connectivity profile with significant projections to 57 and 54 targets for the left and right hemisphere respectively. Connections were mainly to limbic regions such as the orbitofrontal, temporal, anterior cingulate areas, as well as the amygdala and hippocampus. Taken together, this pattern of extrinsic connectivity is consistent with the functional roles ascribed to these areas by the tripartite model (Parent and Hazrati, 1993, 1995), in which cortical and subcortical regions associated with sensorimotor, associative, and affective processing connect predominantly with caudal, dorsal and ventral subregions of the striatum, respectively (Draganski et al., 2008).

## 2.7 Transcriptional signatures of the tripartite striatum

Next, we examined whether connectomically-defined subregions of the striatum show distinct transcriptional profiles. Seventeen tissue samples from the AHBA were located within the caudal striatal cluster, 51 were within the dorsal striatal cluster, and 30 were within the ventral striatal cluster. The limited number of tissue samples available in the AHBA dataset precluded analysis of connectivity-based parcellations where *k* > 3.

### 2.7.1 Transcriptomic prediction of striatal organization

To determine whether gene transcription profiles can accurately differentiate connectomically-defined subregions of the striatum, we first trained an SVM classifier on principal components (PCs) of microarray data drawn from the 98 striatal samples in the AHBA (Fig. 1B). Each PC represents an orthogonal dimension of linear covariance across all genes. Thus, PCs that contribute to classification do so based on the correlated expression of genes acting in concert. PCs were used as predictive features in three one vs. all SVMs, one per striatal cluster. The out-of-sample classification accuracies for each striatal cluster are presented in Figure 3. For each striatal cluster, we repeated the one vs. all classifier ten times, each time using a different PC as a training feature (Fig. 3A). Additionally, we repeated this process using cumulative PCs, up to a total of 10 PCs (Fig. 3B).

**Figure 3.**
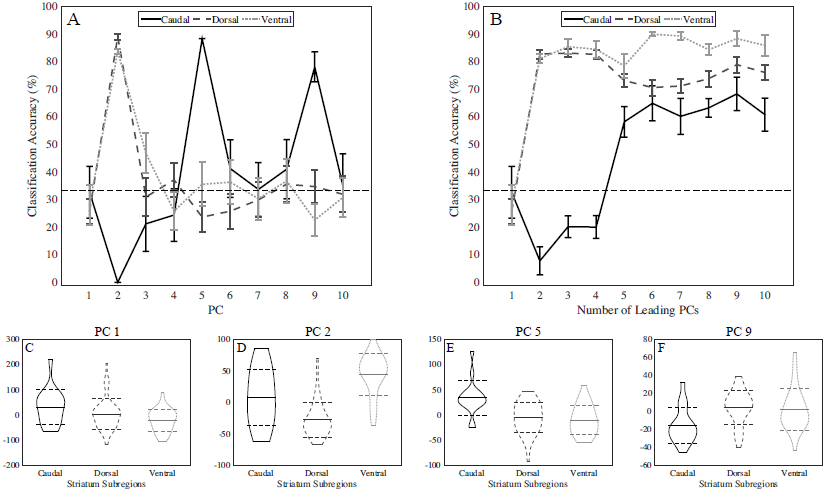
The three connectomically-defined clusters of the striatum have distinct transcriptional profiles. Support vector machine (SVM) classifiers were learnt on principal components (PCs) of transcription data from 98 discrete tissue samples taken from the Allen Human Brain Atlas. SVMs predicted the allocation of tissue samples to each connectomically-defined striatal subregion in a one vs. all framework. Classification accuracy, the proportion of samples in each subregion that are correctly classified, for each striatal subregion is plotted for (A) individual PCs, and (B) as a function of the cumulative number of leading PCs. Accuracy is at approximately chance level for all three striatal clusters when using the first PC. Chance (dashed horizontal lines) was calculated as the mean of the three empirical null distributions collapsed across clusters. (A) Accuracy in-creases sharply when using PC 2 for the dorsal (long-dash line) and ventral (short-dash line) striatal subregions. Accuracy for the caudal subregion (solid line) increases sharply when using PC 5 and again using PC 9. (B) Accuracy is not improved by adding more PCs beyond the first two for the dorsal and ventral subregions. For the caudal subregion, the accuracy of any classifier using a combination of PCs is never as high as the accuracy achieved when just using PC 5 or PC 9. (C-F) Violin plots showing the distribution of PC scores for selected PCs in each of the three striatal subregions. The caudal, dorsal, and ventral striatal subregions contain 17, 51, and 30 tissue samples, respectively. The mean (solid horizontal lines) plus and minus the standard deviation (dashed horizontal lines) of each distribution is also annotated.

For all three striatal clusters, classification accuracy was below chance levels for PC 1, which accounted for 19% of the variance in gene expression across the striatum. This result suggests that the dominant profile of transcriptional variance across the striatum is not predictive of the functional subregions of the tripartite striatum; in other words, it represents transcriptional variability that is common to all three divisions (Fig. 3C). Using PC 2, classification accuracy for the dorsal and ventral striatal clusters increased to approximately 85%, which was significantly greater than chance according to permutation tests (*p*<0.05), suggesting that PC 2, which accounts 11% of the variance in striatal gene expression, is able to clearly separate the dorsal and ventral striatal subregions (Fig. 3D). Classification accuracy for the caudal cluster increases to approximately 90% (*p*<0.05) when using PC 5, and again to approximately 80% (*p*<0.05) when using PC 9, suggesting that a different pattern of gene expression is responsible for separating the caudal cluster from the dorsal and ventral clusters (Fig. 3E–F). Figure 3B reveals that the accuracy of classifying dorsal and ventral subregions does not improve when using more than two PCs. Also, accuracy is hindered when using the first 5 PCs compared to using PC 5 on its own, suggesting that, in this particular application, multi-feature classifiers are less accurate than univariate ones.

### 2.7.2 Transcriptional signatures of striatal subregions

Having established that transcription profiles can be used to robustly classify subregions of the striatum defined by structural connectivity, we wanted to determine whether particular functional groups of genes were driving these results. We focused our analysis on understanding the patterns of gene expression that distinguished the tripartite striatal subregions: PCs 2, 5 and 9, as well as the PC that explained the most variance in the data, PC 1. For each PC, we performed enrichment analysis on the magnitude of the weighting of each individual gene to that PC, allowing us to identify whether genes that weight highly onto these discriminative PCs were associated with any particular GO biological process categories. PCs 1, 2, 5, and 9 were significantly associated with 190, 13, 8, and 47 biological processes in GO, respectively (*p*<0.05, FDR-corrected).

Of the 190 and 47 significant GO categories for PCs 1 and 9, respectively, 115 (60%) and 28 (60%) were nested below the metabolic processes category in the GO hierarchy. Thus, to facilitate interpretation, we reduced the list of significant GO categories for PCs 1 and 9 by taking the top 10 indispensable GO categories as identified using REVIGO (see Methods). GO categories enriched in genes that load strongly onto each PC (1, 2, 5, and 9), including the reduced set for PCs 1 and 9, are displayed in Table 1. PC 1 includes contributions from genes involved in basic cellular functions, including antigen processing, protein localization and tRNA function, and metabolic processes. PC 2 has major contributions from catecholamine-related processes, including genes regulating dopamine receptor signaling and response to amphetamine. PC 5 has contributions from genes involved in synaptic signaling and glutamate secretion and PC 9 includes contributions from genes involved in oxidative metabolism.

**Table 1.**
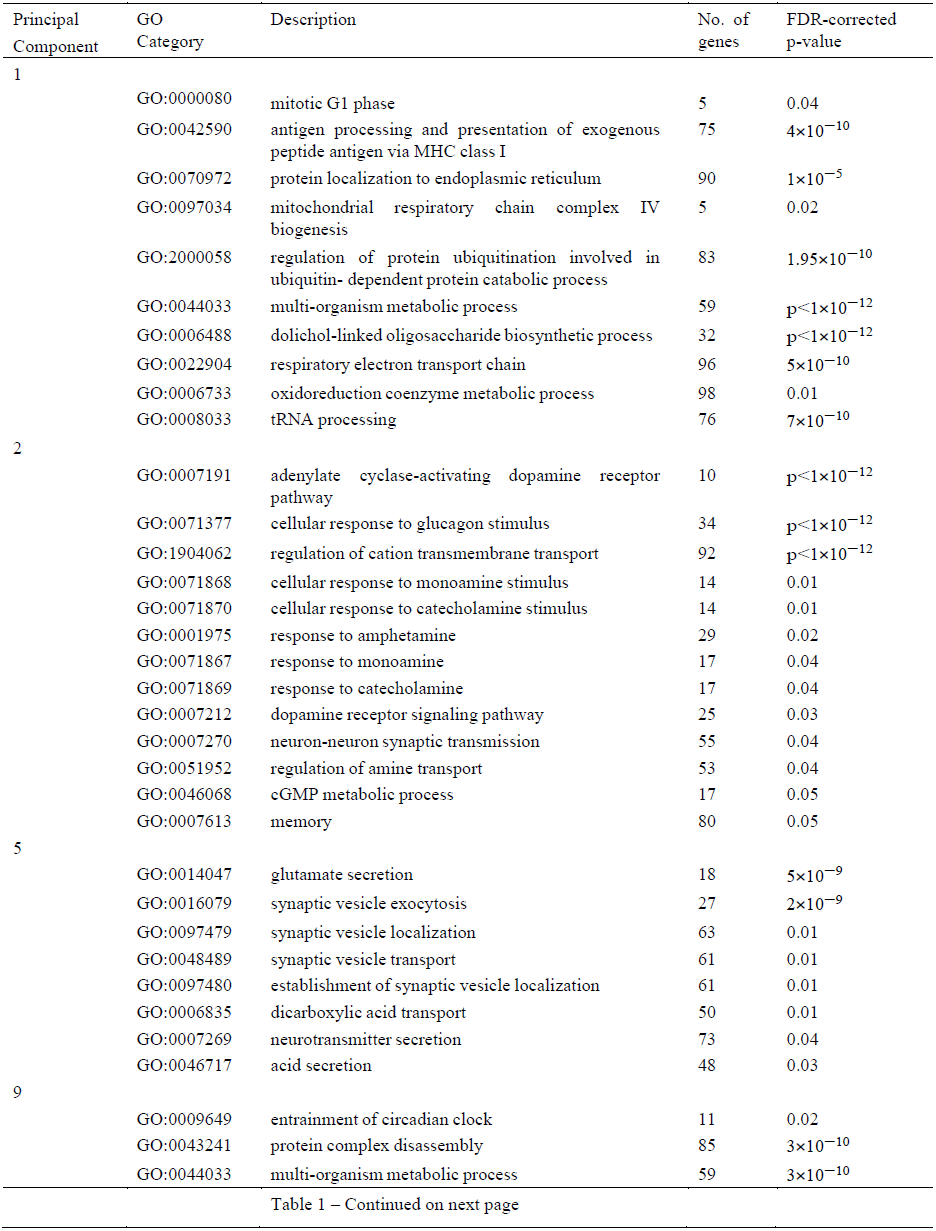
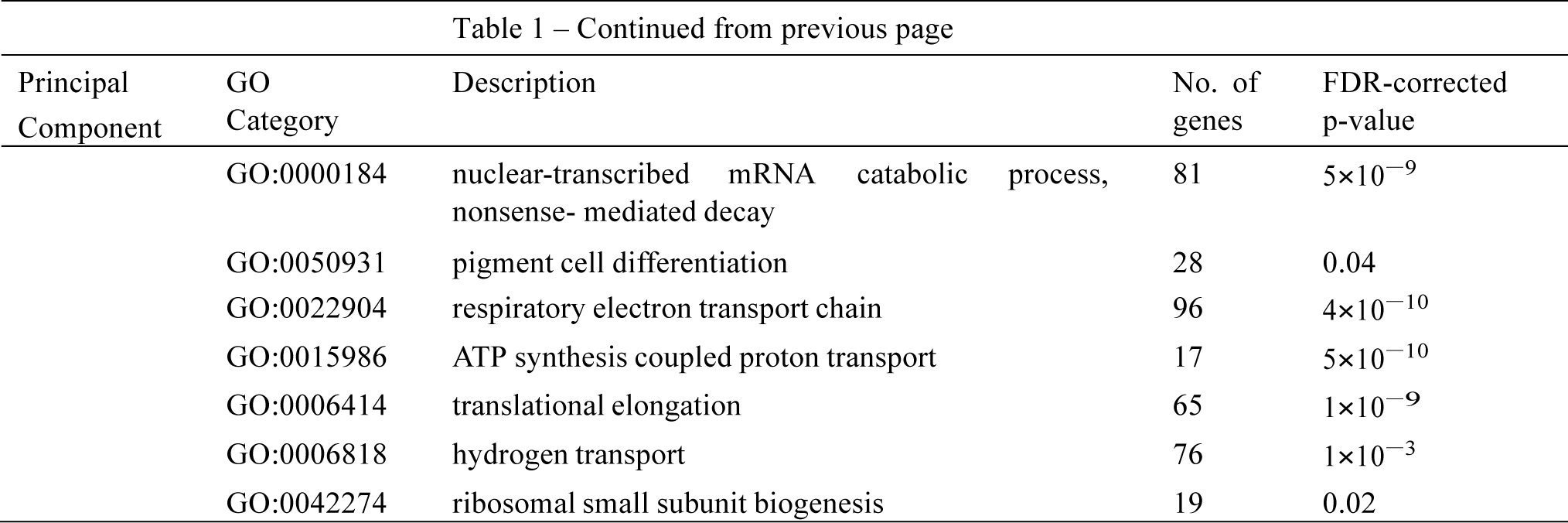
Gene Ontology (GO) biological processes enriched in principle components of transcription data that accurately discriminate between connectomically-defined subregions of the striatum: PC 1 accounts for the largest portion of transcriptional variance but does not classify subregions, PC 2 distinguishes dorsal and ventral subregions, PCs 5 and 9 distinguish the caudal subregion. Significant GO categories are displayed for each PC. GO identification numbers, category names, as well as the number of genes annotated to each category are displayed alongside j3-values from GSR analysis. See text for details.

| Principal Component | GO Category | Description | No. of genes | FDR-corrected p-value |
| --- | --- | --- | --- | --- |
| 1 | GO:0000080 | mitotic G1 phase | 5 | 0.04 |
| | GO:0042590 | antigen processing and presentation of exogenous peptide antigen via MHC class I | 75 | $4 \times 10^{-10}$ |
| | GO:0070972 | protein localization to endoplasmic reticulum | 90 | $1 \times 10^{-5}$ |
|  | GO:0097034 | mitochondrial respiratory chain complex IV biogenesis | 5 | 0.02 |
| | GO:2000058 | regulation of protein ubiquitination involved in ubiquitin- dependent protein catabolic process | 83 | $1.95 \times 10^{-10}$ |
| | GO:0044033 | multi-organism metabolic process | 59 | $p < 1 \times 10^{-12}$ |
| | GO:0006488 | dolichol-linked oligosaccharide biosynthetic process | 32 | $p < 1 \times 10^{-12}$ |
| | GO:0022904 | respiratory electron transport chain | 96 | $5 \times 10^{-10}$ |
|  | GO:0006733 | oxidoreduction coenzyme metabolic process | 98 | 0.01 |
| | GO:0008033 | tRNA processing | 76 | $7 \times 10^{-10}$ |
| 2 | GO:0007191 | adenylate cyclase-activating dopamine receptor pathway | 10 | $p < 1 \times 10^{-12}$ |
| | GO:0071377 | cellular response to glucagon stimulus | 34 | $p < 1 \times 10^{-12}$ |
| | GO:1904062 | regulation of cation transmembrane transport | 92 | $p < 1 \times 10^{-12}$ |
|  | GO:0071868 | cellular response to monoamine stimulus | 14 | 0.01 |
|  | GO:0071870 | cellular response to catecholamine stimulus | 14 | 0.01 |
|  | GO:0001975 | response to amphetamine | 29 | 0.02 |
|  | GO:0071867 | response to monoamine | 17 | 0.04 |
|  | GO:0071869 | response to catecholamine | 17 | 0.04 |
|  | GO:0007212 | dopamine receptor signaling pathway | 25 | 0.03 |
|  | GO:0007270 | neuron-neuron synaptic transmission | 55 | 0.04 |
|  | GO:0051952 | regulation of amine transport | 53 | 0.04 |
|  | GO:0046068 | cGMP metabolic process | 17 | 0.05 |
|  | GO:0007613 | memory | 80 | 0.05 |
| 5 | GO:0014047 | glutamate secretion | 18 | $5 \times 10^{-9}$ |
| | GO:0016079 | synaptic vesicle exocytosis | 27 | $2 \times 10^{-9}$ |
|  | GO:0097479 | synaptic vesicle localization | 63 | 0.01 |
|  | GO:0048489 | synaptic vesicle transport | 61 | 0.01 |
|  | GO:0097480 | establishment of synaptic vesicle localization | 61 | 0.01 |
|  | GO:0006835 | dicarboxylic acid transport | 50 | 0.01 |
|  | GO:0007269 | neurotransmitter secretion | 73 | 0.04 |
|  | GO:0046717 | acid secretion | 48 | 0.03 |
| 9 | GO:0009649 | entrainment of circadian clock | 11 | 0.02 |
| | GO:0043241 | protein complex disassembly | 85 | $3 \times 10^{-10}$ |
| | GO:0044033 | multi-organism metabolic process | 59 | $3 \times 10^{-10}$ |

Table 1 – Continued from previous page
| Principal Component | GO Category | Description | No. of genes | FDR-corrected p-value |
| --- | --- | --- | --- | --- |
| | GO:0000184 | nuclear-transcribed mRNA catabolic process, nonsense-mediated decay | 81 | $5 \times 10^{-9}$ |
|  | GO:0050931 | pigment cell differentiation | 28 | 0.04 |
| | GO:0022904 | respiratory electron transport chain | 96 | $4 \times 10^{-10}$ |
| | GO:0015986 | ATP synthesis coupled proton transport | 17 | $5 \times 10^{-10}$ |
| | GO:0006414 | translational elongation | 65 | $1 \times 10^{-9}$ |
| | GO:0006818 | hydrogen transport | 76 | $1 \times 10^{-3}$ |
|  | GO:0042274 | ribosomal small subunit biogenesis | 19 | 0.02 |

### 2.7.3 Individual variability and genomic predictions of striatal organization

Our initial genomic analysis collapsed tissue samples from six different brains of diverse age, gender, and ethnicity. To examine the robustness of our findings to individual variability in donor brains, we conducted a separate set of individual-level classification analyses using a “leave one brain out” approach (see Methods). Tissue samples from each brain were removed and used as testing data and the remaining five brains were used to train the classifier. Five cumulative PCs in the group-level analysis was the earliest point at which all three SVMs performed greater than chance (Fig. 3B), suggesting that the variance in gene expression measures summarized by the first five PCs is enough to distinguish the caudal, dorsal, and ventral subregions. We therefore used the first five PCs as SVM features in our individual-level analyses. Note that classification accuracy did not vary substantially when using more than the first five PCs. Classification accuracy is displayed for each brain alongside tissue sample cluster coverage and donor demographics in Table 2. Using permutation testing, all six brains showed classification accuracies that were greater than would be expected by chance (*p*<0.05, FDR corrected). High accuracy is driven predominantly by the dorsal and ventral clusters, which may be due to the caudal cluster being less distinctive in its transcriptional signature compared to the others or because there are relatively fewer tissue samples in the caudal cluster when the data from different brains are not pooled. Nonetheless, this analysis supports the robustness of our approach to individual variability in transcriptional activity, and suggests that the ability to predict striatal subdivisions using patterns of gene expression is not driven by age, gender or ethnicity.

**Table 2.** Individual-level genomic prediction of connecomically-defined striatal subdivisions. The Sample Classification heading displays how many tissue samples were accurately classified relative to the total number of tissue samples within each striatal subregion for a given brain. Overall classification accuracy is calculated as the proportion of correctly classified tissue samples for each donor brain across all three subregions. All accuracy results were significant at *p*<0.05, FDR corrected.

| Donor brain | Demographics |  |  | Sample classification<br>(Correct/Total) |  |  | Overall Classification<br>Accuracy |
| --- | --- | --- | --- | --- | --- | --- | --- |
|  | Gender | Ethnicity | Age | Caudal | Dorsal | Ventral |  |
| 1 | Male | African American | 24 | 0/2 | 12/13 | 2/2 | 82% |
| 2 | Male | African American | 39 | 1/4 | 8/8 | 0/1 | 69% |
| 3 | Male | Caucasian | 57 | 0/0 | 5/9 | 6/10 | 58% |
| 4 | Male | Caucasian | 31 | 3/6 | 6/6 | 4/4 | 81% |
| 5 | Female | Hispanic | 49 | 0/1 | 6/7 | 4/4 | 83% |
| 6 | Male | Caucasian | 55 | 2/4 | 6/8 | 7/9 | 71% |

### 2.7.4 Subregional genomic signatures and disease

Having identified which functional groups of genes contribute to separating the three striatal subregions, we next investigated the overlap between the sets of genes that correlate with PCs and genes implicated in risk for different diseases. We focused our analysis on PCs 2, 5, and 9. For each PC, we took the top 10% of genes with the strongest absolute correlations to PCs, as a gene set (*n* = 1 934 genes). Next, we used Autworks to generate a list of diseases that overlapped significantly with each PC’s gene set. The top 10% of genes correlating with PCs 2, 5, and 9 overlapped significantly with genes associated with 6, 12, and 1 different diseases, respectively (*p*<0.05, FDR-corrected). These results are listed in Table 3. Genes correlated with PC 9 overlapped with general disease annotations relating to metabolic brain disorders and mitochondrial disease. Genes correlated with PC 2 were over-represented in psychotic disorders, including schizophrenia and affective psychosis, consistent with the strong involvement of catecholamine (particularly dopaminergic) genes (Table 1). It is also consistent with the accurate classification of the dorsal and ventral subregions by PC2, given that the former is heavily implicated in psychosis (Fornito et al., 2013, Howes et al., 2009). Genes correlated with PC 5 overlapped with genes implicated in schizophrenia, epilepsy and other developmental disorders such as autism and intellectual disability, consistent with these genes playing a role in general synaptic function and glutamate transmission (Table 1).

**Table 3.** Disease associations for principle components (PCs) of transcription data collected from 98 tissue samples spanning the striatum. Each PC represents variation in transcription data that distinguishes between connectomically-defined subregions of the striatum. For each PC, gene expression data was correlated with PC scores and the top 10% were used with Autworks to determine disease associations. All results were significant at *p*<0.05 (permutation test, FDR-corrected).

| Principle component | Associated diseases |
| --- | --- |
| 2 | Schizophrenia and Disorders with Psychotic Features<br>Schizophrenia<br>Mental Disorders<br>Nervous System Diseases<br>Bipolar Disorder<br>Affective Disorders, Psychotic |
| 5 | Mental Disorders Diagnosed in Childhood<br>Mental Disorders Epilepsy, Generalized Epilepsy<br>Epilepsy, Absence<br>Schizophrenia<br>Schizophrenia and Disorders with Psychotic Features<br>Neurobehavioral Manifestations<br>Child Development Disorders, Pervasive<br>Intellectual Disability Autistic Disorder<br>Neurodegenerative Diseases |
| 9 | Mitochondrial Diseases |

To test the robustness of our findings, we re-ran the analysis selecting genes that were significantly correlated to each PC (*p*<0.05, FDR-corrected), rather than simply using the top 10%. Similar overall findings were obtained, but with some additional disease associations for some PCs, likely because the number of genes assigned to each PC was free to vary in this analysis (see Supplementary Table 1). For example, genes significantly correlated with PC 2 were also associated with X-linked mental retardation. Interestingly, the list of genes significantly correlating with PC 5 also included movement disorders and Parkinson’s disease. Given that PC 5 distinguished the caudal from dorsal and ventral striatum, this finding is consistent with evidence that pathology of the caudal striatum may be more prominent in Parkinson’s disease (Kish et al., 1988).

### 2.7.5 Validation against other striatal parcellations

Having analyzed the transcriptional signatures of striatal subregions defined using k-means clustering of connectomic data, we next investigated whether we could obtain comparable classification accuracy when tissue samples were assigned to striatal subregions defined using different criteria. We considered two alternative striatal parcellations: one that used anatomical boundaries (Desikan et al., 2006) and one that defined a tripartite division using an alternative connectomic strategy, which used winner-takes-all assignment of striatal voxels based on their connectivity to cortical lobes (Tziortzi et al., 2014) (see section 2.5.5).

The anatomical parcellation was based on the same initial striatal mask as our own parcellation, so the spatial coverage of the AHBA that the anatomical atlas provides is identical to ours, as are the principal components of genetic variance. The only difference is the subregion label assigned to each tissue sample, with the anatomical parcellation assigning 26 tissue samples to the caudate, 57 to the putamen and 15 to the nucleus accumbens. Classification accuracy for the caudate and ventral striatum showed greater than chance classification accuracy, but only the caudate was significantly different from chance at *p*<0.05 (permutation test, FDR-corrected; Fig S1A). Accuracy for the caudate subregion was driven by PC 2, which is the same PC that separated the dorsal and ventral subregions in our parcellation. Accuracy for the putamen was never significantly greater than chance (according to a permutation test) for any of the top 10 PCs.

The alternative connectomic parcellation, developed by Tziortzi et al., (2014), is derived from the Harvard-Oxford atlas, which is based on a smaller mask of the striatum than the Desikan-Killiany atlas. As a result, only 63 of the original 98 tissue samples could be assigned to a striatal subregion in the Tziortzi parcellation; 23 in the caudal subregion, 30 in the dorsal subregion, and 10 in the ventral subregion. To ensure direct comparability to the analyses using the other parcellations, we assigned the remaining 35 tissue samples to the nearest subregion of the Tziortzi parcellation using the same Euclidean distance method outline in section 2.5. This resulted in 32 samples being located in the caudal subregion, 36 in the dorsal subregion, and 30 in the ventral subregion. Classification accuracy for the dorsal and ventral subregions reached similarly high levels of classification accuracy compared our own parcellation (Fig S1B), but only accuracy for the dorsal subregion was significantly different from chance (*p*<0.05, FDR-corrected) on PC 2. Accuracy for the caudal subregion was never significantly greater than chance for any of the top 10 PCs. Thus, unlike in our parcellation, neither of the two alternative striatal parcellations examined here were able to classify all three subregions with accuracy significantly greater than chance.

## 3 Discussion

We sought to determine whether connectomically-defined regions of the brain carry distinctive transcriptional signatures. We focused in particular on the human striatum, a region that has a well-characterized functional and chemical organization and which has been implicated in a diverse range of human behaviors and disorders. We demonstrated that the human striatum can be delineated into distinct subregions, based on extrinsic connectivity to the rest of the brain, and that these subregions can be distinguished with high accuracy based on their gene expression profile. In support of previous literature (Levey et al., 1993, Gurevich, 1999), our results show that dopamine receptor signaling and response to amphetamine transcripts are among the dominant sources of genetic variation separating the dorsal and ventral subregions of the striatum, while transcripts associated with glutamate secretion and metabolic processes separate the caudal subregion. These subregional genetic signatures were also linked to several disorders that are thought to be associated with the striatum, such as schizophrenia, bipolar, and autism. The multi-modal methodology developed in this study provides a framework for future work combining brain connectivity and gene expression data to provide a more complete understanding of brain organization, both in terms of macroscopic axonal organization and molecular architecture.

### 3.1 Connectomic parcellation of the striatum

We used unbiased, data-driven clustering analysis of extrinsic connectivity to replicate the tripartite partition of the striatum (Parent and Hazrati, 1995, 1993, Parent, 1990). We found that striatal subregions showed expected patterns of whole-brain connectivity: the caudal cluster was strongly connected with sensorimotor cortex, the dorsal cluster with prefrontal areas, and the ventral cluster with limbic regions. Our method extends previous attempts at parcellating the striatum based on structural connectivity in two key ways. First, we used a large number of connectivity targets compared to previous attempts that only used a small number of cortical lobe targets (Tziortzi et al., 2014, Bohanna et al., 2011), reducing the heterogeneity within targets and enhancing variability of the connectivity profiles of individual striatal voxels. Second, rather than threshold and binarise connectivity profiles (e.g., Draganski et al., 2008) or just take the target with the highest probability (e.g., Tziortzi et al., 2014), we clustered the connectivity profiles across all targets. By taking account of a broader array of data, our approach subjected the tripartite hypothesis to a stringent test.

At higher levels of granularity, we found evidence for a rostrocaudal gradient, consistent with past diffusion MRI (Draganski et al., 2008) research in humans. This gradient is likely to support functionally specialized circuits within the broader ventral-affective, dorsal-associative and caudal-sensorimotor partitions. There were no abrupt changes in regional boundaries as a function of increasing k, indicating that these finer-grained divisions were nested within the ventral, dorsal and caudal clusters, consistent with a hierarchical organization of extrinsic connectivity in the striatum.

An important difference between our parcellation and those of previous attempts (e.g., Tziortzi et al., 2014, Bohanna et al., 2011) is that, in addition to the rostrocaudal and dorsoventral divisions, we also found that subregions split along a medial-lateral axis. A common view is that the distinction between the medial and lateral parts of the striatum, which corresponds to the anatomical distinction between the caudate and putamen in humans, functionally subserves goal-directed and habitual behavior, respectively (Haruno and Kawato, 2006, Jog, 1999, Balleine et al., 2007). However, single-neuron electrophysiology in rats has shown that goal-driven and habitual behaviors are encoded for across the medial and lateral compartments of the caudate-putamen (Stalnaker et al., 2010), suggesting that the gross medial-lateral distinction may be too simplistic to capture differences in function. Indeed, nonhuman primate research suggests that the ability to carry out habitual motor sequences is impaired when activity in the lateral section of the putamen is chemically suppressed, whereas suppressing the medial section of the putamen results in impaired acquisition of novel motor sequences (Miyachi et al., 1997). The medial-lateral gradients within the caudate and putamen observed here are also consistent with the existence of a dorsocentral region of the striatum, previously identified in rodent tract-tracing studies (Reep and Corwin, 1999, Reep et al., 1987). This region is a major site of corticostriatal projections from the agranular cortex (Reep et al., 1987), which is thought to be a homologue to primate area 8, a frontal region linked to executive processing (Reep and Corwin, 1999). Similar medial-lateral gradients have been observed in human DWI (Draganski et al., 2008) and resting-state functional MRI (Jung et al., 2014) data.

### 3.2 The transcriptional signature of the tripartite striatum

Our findings indicate that variation in transcriptional measures taken from 19 343 genes in 98 samples spanning the extent of the striatum can accurately predict connectomically-defined subregions of the tripartite striatum. Accurate prediction was retained at the level of individual brains, despite differences in demographics that can influence gene expression, such as ethnicity (Jorde and Wooding, 2004), demonstrating a strong link between connectomically-defined subregions of the striatum and gene expression that is conserved across individuals. Classification accuracy was higher for our connectivity-based parcellation, which incorporated multivariate voxel connectivity profiles, compared to parcellations based on anatomical boundaries (Desikan et al., 2006) or coarser winner-takes-all approaches (Tziortzi et al., 2014). These results suggest that connectomic parcellations can be used to recover meaningful functional subdivisions of the brain, and that incorporating rich, multivariate profiles of interregional connectivity may enhance the biological validity of these divisions.

We examined enrichment of principal components of transcriptional variance across subregion of the striatum. The first principal component of transcription data (PC 1), which captured the dominant trend of the transcriptional variation of all genes across striatal tissue samples, did not differentiate the ventral, dorsal and caudal divisions. Enrichment analysis revealed processes that are fundamental to cellular function and signaling, such as protein transport, localization and ubiquitination, antigen processing, tRNA processing and metabolic processes that are integral for supplying the energy required for neuronal activity (Attwell and Laughlin, 2001). The first principal component is thus dominated by genes related to essential processes related to cellular function in general, rather than specific properties that may differentiate distinct striatal subdivisions.

The second PC significantly discriminated between dorsal and ventral subregions of the striatum. We found that PC 2 showed enrichment for biological processes such as dopamine receptor signaling and response to amphetamines, monoamines, and catecholamines. These categories included genes coding for dopamine receptors in the D1, D2, and D3 families as well as cannabinoid receptors. The dorsal and ventral subregions are known to differ in terms of dopamine receptor density (Gurevich, 1999, Levey et al., 1993) and have been heavily implicated in the dopaminergic dysfunction that is characteristic of schizophrenia (Dandash, Pantelis, & Fornito, 2016; Fornito et al., 2013, Howes et al., 2009). For example, dopamine D2 receptors, which are expressed more in the dorsal striatum relative to ventral (Levey et al., 1993, Olsen et al., 2008), are the primary target for most antipsychotics used to treat the positive symptoms of schizophrenia (Seeman and Lee, 1975). Conversely, dopamine D3 receptors, which more strongly expressed in the ventral subregion of the striatum (Gurevich, 1999), may be linked to the negative symptoms of schizophrenia (Simpson et al., 2014). Endogenous signaling at cannabinoid receptors can modulate glutamatergic and dopaminergic signaling in the striatum (Robbe et al., 2003), and through this mechanism, the endocannabinoid system has been implicated in risk for disorders associated with striatal dysfunction, such as Parkinson’s disease, Huntington’s disease, drug addiction and schizophrenia (van der Stelt and Di Marzo, 2003). It is thus not surprising that genes loading onto PC2 were also associated with risk for schizophrenia.

The enrichment of the ‘response to amphetamines’ GO category may reflect the segregation of dopaminergic projections from the midbrain within the striatum. Mesolimbic projections from the ventral tegmental area terminate predominantly in the ventral striatum whereas nigrostriatal projections from the substantia nigra terminate predominantly in the dorsal striatum (Haber et al., 2000). Research into acute amphetamine exposure in non-users has demonstrated a greater reduction in dopamine D2 receptor availability in the ventral striatum compared to the dorsal striatum (Martinez et al., 2003), suggesting that the mesolimbic pathway is more responsive to amphetamines compared to the nigrostriatal pathway.

PCs 5 and 9 accurately classified the caudal subregion of the striatum. Our enrichment analyses revealed that the top GO category for PC 5 was ‘glutamate secretion’. A notable characteristic of our parcellation is that the caudate and the putamen are both divided into medial and lateral segments (Fig. 2A), with the lateral segments of each appearing in the caudal subregion and the medial segments appearing in the dorsal subregion. Consistent with this characteristic, autoradiography in non-human primates has shown that class-II metabotropic glutamate receptors, genes for which are annotated to the ‘glutamate secretion’ category (e.g., GRM2), are characterized by an increasing ventrolateral-to-dorsomedial concentration gradient in both the caudate and the putamen (Beveridge et al., 2011). This suggests that our connectivity-based tripartite parcellation captures functional differences (i.e., sensorimotor vs. associative processing) within the caudate and putamen that may be subserved by differences in glutamate concentrations.

### 3.3 Limitations and conclusions

The potential for crossing fibers to pass through a single voxel can cause problems for diffusion tractography algorithms. We reconstructed fiber tracts using a ball-and-stick crossing fiber model (Behrens et al., 2007), coupled with high-resolution, multi-shell diffusion data which enhances our sensitivity to accurately reconstruct complex fiber trajectories (Sotiropoulos et al., 2013). Nonetheless, diffusion tractography is known to suffer from problems of sensitivity and specificity which can hinder accurate tract reconstruction (Thomas et al., 2014). The limited resolution of diffusion data also means that we are unable to identify small substructures. For example, the nucleus accumbens is known to comprise and core and shell region, but it is not possible to resolve this level of detail with current imaging technologies. Our findings thus represent a parcellation of the macroscale structure of the striatum.

The AHBA offers an unprecedented spatial coverage of gene expression across the entire brain, providing new opportunities to study regional variations in transcriptional activity. The scale and challenge of acquiring these data necessarily means that they are limited in some respects. For example, data are only available from six donor brains, and we had to pool data across these brains to obtain sufficient spatial coverage of the entire striatum. However, our leave-one-brain-out analysis confirmed that our findings are robust to inter-individual variability in the donor brains. Our analysis also only focused on tissue samples taken from the left hemisphere because only left hemisphere samples were available in four of the six donor brains. Prior analysis of these data has shown that hemispheric differences in gene expression are relatively minor (Hawrylycz et al., 2012), but an interesting open question is whether the transcriptional signatures identified here are symmetrically represented across hemispheres.

Despite combining data across the six donor brains in the AHBA, low tissue sample density within the striatum still precluded genetic analysis of striatal parcellations with more than three subregions. This limitation is pertinent given suggestions that hard segmentation parcellation methods oversimplify the anatomical organization of brain regions (see Lambert et al., 2012, Alkemade and Forstmann, 2014, Lambert et al., 2015, for a discussion) and that mapping connectivity gradients may be a more accurate model of anatomical organization (e.g., Draganski et al., 2008). Denser spatial sampling of gene expression in the striatum would allow for more fine grained investigation of the striatum and other brain regions, but at the spatial resolution currently available a relatively coarse tripartite parcellation was necessary to examine differences in gene transcription. An alternative approach would be to perform clustering on the gene expression data rather than the diffusion data. However, with relatively few samples in the tail of caudate, obtaining a robust genetic signature for this region is problematic without denser spatial sampling of gene transcription. Our approach took advantage of the complete spatial coverage of the striatum by diffusion MRI, while also offering a means to identify distinct transcriptional signatures of each subregion. Sufficient tissue sample coverage at an individual level would also remove the need to pool the AHBA data, thus removing the caveat of inter-individual variability, although our leave-one-brain-out analysis suggests that our results are robust to such variability.

In summary, we showed that variation in gene expression spanning the whole human genome could accurately distinguish between connectomically-defined striatal subregions. These results were driven by genes related to dopamine and glutamate signaling and neuronal communication, and were also implicated in risk for a range of different brain disorders associated with striatal dysfunction. Our methodology offers a general framework for the crossvalidation of connectomic parcellations with genetic data.

## Acknowledgements

The authors thank Dr. Orwa Dandash for helpful feedback on the manuscript. L.P. was supported by an Australian Postgraduate Award. B.F. was supported by a National Health and Medical Research Council Early Career Fellowship (ID: 1089718). M.Y. was supported by a National Health and Medical Research Council Fellowship (ID: APP1021973), Monash University and the David Winston Turner Endowment Fund. A.F. was supported by an Australian Research Council Future Fellowship (ID: FT130100589) and National Health and Medical Research Council Project grants (ID: 3251213 and 3251250). Data were provided [in part] by the Human Connectome Project, WU-Minn Consortium (Principal Investigators: David Van Essen and Kamil Ugurbil; 1U54MH091657) funded by the 16 NIH Institutes and Centers that support the NIH Blueprint for Neuroscience Research; and by the McDonnell Center for Systems Neuroscience at Washington University.

---

**Supplementary Figure 1.**
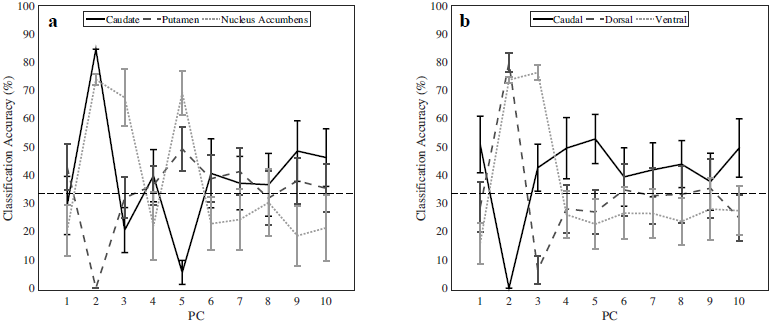
Replication of support vector machine (SVM) classifiers using (*A*) an anatomical parcellation as well as (*B*) an alternative tripartite parcellation to separate tissue samples taken from the Allen Human Brain Atlas (AHBA). SVMs predicted the allocation of AHBA tissue sample to each striatal subregion in a one vs. all framework. Classification accuracy, the proportion of samples in each subregion that are correctly classified, for each striatal subregion is plotted for individual PCs. In both parcellations, classification accuracy greater than chance is never achieved for the dorsal subregion or the putamen. Note that PCs are not comparable between parcellations as they likely represent different sources of variance (see main text).

**Table S1.**
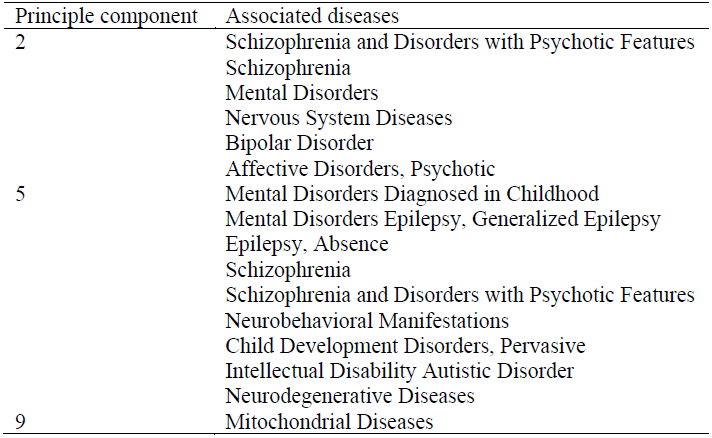
Disease associations for principle components (PCs) of transcription data collected from 98 tissue samples spanning the striatum. Each PC represents variation in transcription data that distinguishes between connectomically-defined subregions of the striatum. For each PC, gene expression data was correlated with PC scores and the correlations significant under permutation tests (see main text) were used with Autworks to determine disease associations. All results were significant at *p*<0.05 (permutation test, FDR-corrected).

| Principle component | Associated diseases |
| --- | --- |
| 2 | Schizophrenia and Disorders with Psychotic Features<br>Schizophrenia<br>Mental Disorders<br>Nervous System Diseases<br>Bipolar Disorder<br>Affective Disorders, Psychotic<br>X-linked Mental Retardation |
| 5 | Mental Disorders Diagnosed in Childhood<br>Mental Disorders<br>Epilepsy, Generalized<br>Epilepsy<br>Epilepsy, Absence<br>Schizophrenia<br>Schizophrenia and Disorders with Psychotic Features<br>Neurobehavioral Manifestations<br>Child Development Disorders, Pervasive<br>Intellectual Disability<br>Autistic Disorder<br>Neurodegenerative Diseases<br>Parkinson Disease, Secondary<br>Movement Disorders<br>Seizures<br>Gait Disorders, Neurologic<br>Mood Disorders |
| 9 | Mitochondrial Diseases |

## References

1. Alcauter, S, Lin, W, Keith Smith, J, Gilmore, JH & Gao, W (2015). Consistent Anterior-Posterior Segregation of the Insula During the First 2 Years of Life. Cereb.Cortex, 25, 1176–1187.

2. Alkemade, A & Forstmann, BU (2014). Do we need to revise the tripartite subdivision hypothesis of the human subthalamic nucleus (STN)? Neuroimage, 95, 326–329.

3. Andersson, JL, Skare, S & Ashburner, J (2003). How to correct susceptibility distortions in spin-echo echo-planar images: application to diffusion tensor imaging. Neuroimage, 20, 870–888.

4. Anwander, A, Tittgemeyer, M, von Cramon, D, Friederici, A & Knosche, T (2006). Connectivity-Based Parcellation of Broca’s Area. Cereb. Cortex, 17, 816–825.

5. Ashburner, M, Ball, Ca, Blake, Ja, Botstein, D, Butler, H, Cherry, JM, Davis, AP, Dolinski, K, Dwight, SS, Eppig, JT, Harris, Ma, Hill, DP, Issel-Tarver, L, Kasarskis, A, Lewis, S, Matese, JC, Richardson, JE, Ringwald, M, Rubin, GM & Sherlock, G (2000). Gene Ontology: tool for the unification of biology. Nat. Genet. 25, 25–29.

6. Attwell, D & Laughlin, SB (2001). An Energy Budget for Signaling in the Grey Matter of the Brain. J. Cereb Blood Flow Metab. 21, 1133–1145.

7. Baliki, MN, Mansour, A, Baria, AT, Huang, L, Berger, SE, Fields, HL & Apkarian, AV (2013). Parceling Human Accumbens into Putative Core and Shell Dissociates Encoding of Values for Reward and Pain. J. Neurosci. 33, 16383–16393.

8. Balleine, BW, Delgado, MR & Hikosaka, O (2007). The Role of the Dorsal Striatum in Reward and Decision-Making. J. Neurosci. 27, 8161–8165.

9. Behrens TEJ & Johansen-Berg H (2005). Relating connectional architecture to grey matter function using diffusion imaging. Philos. Trans. R. Soc. B Biol. Sci. 360, 903–911.

10. Behrens TEJ, Johansen-Berg, H, Woolrich, MW, Smith, SM, Wheeler-Kingshott, CaM, Boulby, Pa, Barker, GJ, Sillery, EL, Sheehan, K, Ciccarelli, O, Thompson, aJ, Brady, JM & Matthews, PM (2003). Non-invasive mapping of connections between human thalamus and cortex using diffusion imaging. Nat. Neurosci. 6, 750–757.

11. Behrens, T, Berg, HJ, Jbabdi, S, Rushworth, M & Woolrich, M (2007). Probabilistic diffusion tractography with multiple fibre orientations: What can we gain? Neuroimage, 34, 144–155.

12. Behrens, T, Woolrich, M, Jenkinson, M, Johansen-Berg, H, Nunes, R, Clare, S, Matthews, P, Brady, J & Smith, S (2003). Characterization and propagation of uncertainty in diffusion-weighted MR imaging. Magn. Reson. Med. 50, 1077–1088.

13. Benjamini, Y & Hochberg, Y (1995). Controlling the false discovery rate: a practical and powerful approach to multiple testing. J. R. Statisitical Soc. 57, 289–300.

14. Beveridge, T, Smith, H, Nader, M & Porrino, L (2011). Group II metabotropic glutamate receptors in the striatum of non-human primates: Dysregulation following chronic cocaine self-administration. Neurosci. Lett. 496, 15–19.

15. Bohanna, I, Georgiou-Karistianis, N & Egan, GF (2011). Connectivity-based segmentation of the striatum in Huntington’s disease: vulnerability of motor pathways. Neurobiol. Dis. 42, 475–81.

16. Dandash, O, Fornito, A, Lee, J, Keefe RSE Chee MWL Adcock, RA, Pantelis, C, Wood, SJ & Harrison, BJ (2014). Altered Striatal Functional Connectivity in Subjects With an At-Risk Mental State for Psychosis. Schizophr. Bull. 40, 904–913.

17. Dandash, O., Pantelis, C., & Fornito, A. (2016). Dopamine, fronto-striato-thalamic circuits and risk for psychosis. Schizophrenia Research. https://doi.org/10.1016/j.schres.2016.08.020

18. DeLong, MR & Wichmann, T (2007). Circuits and Circuit Disorders of the Basal Ganglia. Neurol. Rev. 64.

19. Desikan, RS, S’egonne, F, Fischl, B, Quinn, BT, Dickerson, BC, Blacker, D, Buckner, RL, Dale, AM, Maguire, RP, Hyman, BT, Albert, MS & Killiany, RJ (2006). An automated labeling system for subdividing the human cerebral cortex on MRI scans into gyral based regions of interest. Neuroimage, 31, 968–980.

20. Desikan, R. S., Ségonne, F., Fischl, B., Quinn, B. T., Dickerson, B. C., Blacker, D., … Killiany, R. J. (2006). An automated labeling system for subdividing the human cerebral cortex on MRI scans into gyral based regions of interest. NeuroImage, 31(3), 968–980. https://doi.org/10.1016/j.neuroimage.2006.01.021

21. Draganski, B, Kherif, F, Kloppel, S, Cook, PA, Alexander, DC, Parker GJM, Deichmann, R, Ashburner, J & Frackowiak, RSJ (2008). Evidence for Segregated and Integrative Connectivity Patterns in the Human Basal Ganglia. J. Neurosci. 28, 7143–7152.

22. Eickhoff, SB, Thirion, B, Varoquaux, G & Bzdok, D (2015). Connectivity-based parcellation: Critique and implications. Hum. Brain Mapp. 36, 4771–4792.

23. Everitt, BJ & Robbins, TW (2013). From the ventral to the dorsal striatum: devolving views of their roles in drug addiction. Neurosci. Biobehav. Rev. 37, 1946–54.

24. Fan L, Li H, Zhuo J, Zhang Y, Wang J, Chen L, Yang Z, Chu C, Xie S, Laird AR, Fox PT, Eickhoff SB, Yu C & Jiang, T (2016). The Human Brainnetome Atlas: A New Brain Atlas Based on Connectional Architecture. Cereb. Cortex 26 3508–3526.

25. Fornito A, Harrison BJ, Goodby E, Dean, A, Ooi C, Nathan PJ, Lennox BR, Jones, PB, Suckling J & Bullmore, ET (2013). Functional dysconnectivity of corticostriatal circuitry as a risk phenotype for psychosis. JAMA psychiatry, 70, 1143–51.

26. French L & Pavlidis, P (2011). Relationships between Gene Expression and Brain Wiring in the Adult Rodent Brain. PLoS Comput. Biol. 7, e1001049.

27. Fulcher BD & Fornito A (2016). A transcriptional signature of hub connectivity in the mouse connectome. Proc. Natl. Acad. Sci. I, 201513302.

28. Giménez-Amaya JM, McFarland NR, De Las Heras, S & Haber SN (1995). Organization of thalamic projections to the ventral striatum in the primate. J. Comp. Neurol. 354, 127–149.

29. Glasser MF, Sotiropoulos SN, Wilson JA, Coalson TS, Fischl B, Andersson JL, Xu J, Jbabdi S, Webster M, Polimeni JR, Van Essen DC & Jenkinson M (2013). The minimal preprocessing pipelines for the Human Connectome Project. Neuroimage, 80, 105–124.

30. Gordon EM, Laumann TO, Adeyemo B, Huckins JF, Kelley WM & Petersen SE (2016). Generation and Evaluation of a Cortical Area Parcellation from Resting-State Correlations. Cereb. Cortex, 26, 288–303.

31. Gurevich E (1999). Distribution of Dopamine D3 Receptor Expressing Neurons in the Human Forebrain Comparison with D2 Receptor Expressing Neurons. Neuropsychopharmacology, 20, 60–80.

32. Haber SN, Fudge JL & McFarland NR (2000). Striatonigrostriatal pathways in primates form an ascending spiral from the shell to the dorsolateral striatum. J. Neurosci. 20, 2369–82.

33. Haber SN (2003). The primate basal ganglia: parallel and integrative networks. J. Chem. Neuroanat. 26, 317–330.

34. Harrison BJ, Soriano-Mas, C, Pujol J, Ortiz H, López-Solà M, Hernández-Ribas, R, Deus, J, Alonso, P, Yücel, M, Pantelis, C, Menchon, JM & Cardoner, N (2009). Altered corticostriatal functional connectivity in obsessive-compulsive disorder. Arch. Gen. Psychiatry, 66, 1189–200.

35. Haruno M & Kawato M (2006). Different neural correlates of reward expectation and reward expectation error in the putamen and caudate nucleus during stimulus-action-reward association learning. J. Neurophysiol. 95, 948–59.

36. Hastie, T, Tibshirani, R & Friedman, J (2009). The Elements of Statistical Learning Springer Series in Statistics. Springer New York, New York, NY.

37. Hawrylycz M, Miller JA, Menon V, Feng D, Dolbeare T, Guillozet-Bongaarts AL, Jegga AG, Aronow BJ, Lee CK, Bernard A, Glasser MF, Dierker DL, Menche J, Szafer A, Collman F, Grange P, Berman KA, Mihalas S, Yao Z, Stewart L, Barabási AL, Schulkin J, Phillips J, Ng L, Dang C, Haynor DR, Jones A, Van Essen DC, Koch C & Lein, E (2015). Canonical genetic signatures of the adult human brain. Nat. Neurosci. 18, 1832–44.

38. Hawrylycz MJ, Lein ES, Guillozet-Bongaarts AL, Shen EH, Ng L, Miller JA, van de Lagemaat LN, Smith KA, Ebbert A, Riley ZL, Abajian C, Beckmann CF, Bernard A, Bertagnolli D, Boe AF, Cartagena PM, Chakravarty MM, Chapin M, Chong J, Dalley RA, Daly BD, Dang C, Datta S, Dee N, Dolbeare TA, Faber V, Feng D, Fowler DR, Goldy J, Gregor BW, Haradon Z, Haynor DR, Hohmann JG, Horvath S, Howard RE, Jeromin A, Jochim JM, Kinnunen M, Lau C, Lazarz ET, Lee C, Lemon TA, Li, L, Li Y, Morris JA, Overly CC, Parker PD, Parry SE, Reding M, Royall JJ, Schulkin J, Sequeira PA, Slaughterbeck CR, Smith SC, Sodt AJ, Sunkin SM, Swanson BE, Vawter MP, Williams D, Wohnoutka P, Zielke HR, Geschwind DH, Hof PR, Smith SM, Koch C, Grant SGN & Jones AR (2012). An anatomically comprehensive atlas of the adult human brain transcriptome. Nature, 489, 391–399.

39. Hawrylycz M, Bernard A, Lau C, Sunkin SM, Chakravarty MM, Lein ES, Jones AR & Ng L (2010). Areal and laminar differentiation in the mouse neocortex using large scale gene expression data. Methods, 50, 113–21.

39 Howes OD, Montgomery AJ, Asselin MC, Murray RM, Valli I, Tabraham P, Bramon-Bosch E, Valmaggia L, Johns L, Broome M, McGuire PK & Grasby PM (2009). Elevated Striatal Dopamine Function Linked to Prodromal Signs of Schizophrenia. Arch. Gen. Psychiatry, 66, 13.

40 Jbabdi S, Sotiropoulos SN, Savio AM, Graña M & Behrens TEJ (2012). Model-based analysis of multishell diffusion MR data for tractography: how to get over fitting problems. Magn. Reson. Med. 68, 1846–55.

41 Jog MS (1999). Building Neural Representations of Habits. Science, 286, 1745–1749.

42 Johansen-Berg H, Behrens TEJ, Robson MD, Drobnjak I, Rushworth MFS, Brady JM, Smith SM, Higham DJ & Matthews PM (2004). Changes in connectivity profiles define functionally distinct regions in human medial frontal cortex. Proc. Natl. Acad. Sci. 101, 13335–13340.

43 Jorde LB & Wooding SP (2004). Genetic variation, classification and ‘race’. Nat. Genet. 36, S28–S33.

44 Jung WH, Jang JH, Park JW, Kim E, Goo EH, Im OS & Kwon JS (2014). Unravelling the Intrinsic Functional Organization of the Human Striatum: A Parcellation and Connectivity Study Based on Resting-State fMRI. PLoS One, 9, e106768.

45 Kaufman A, Dror, G, Meilijson I & Ruppin E (2006). Gene expression of Caenorhabditis elegans neurons carries information on their synaptic connectivity. PLoS Comput. Biol. 2, e167.

46 Kish SJ, Shannak K & Hornykiewicz O (1988). Uneven pattern of dopamine loss in the striatum of patients with idiopathic Parkinson’s disease. Pathophysiologic and clinical implications. N. Engl. J. Med. 318, 876–80.

47 Koehler S, Ovadia-Caro S, van der Meer E, Villringer A, Heinz A, Romanczuk-Seiferth N & Margulies DS (2013). Increased functional connectivity between prefrontal cortex and reward system in pathological gambling. PLoS One, 8, e84565.

48 Krienen FM, Yeo BTT, Ge T, Buckner RL & Sherwood CC (2016). Transcriptional profiles of supragranular enriched genes associate with corticocortical network architecture in the human brain. Proc. Natl. Acad. Sci. 113, E469–E478.

49 Lambert C, Zrinzo L, Nagy Z, Lutti A, Hariz M, Foltynie T, Draganski B, Ashburner J & Frackowiak R (2012). Confirmation of functional zones within the human subthalamic nucleus: Patterns of connectivity and subparcellation using diffusion weighted imaging. Neuroimage, 60, 83–94.

50 Lambert C, Zrinzo L, Nagy Z, Lutti A, Hariz M, Foltynie T, Draganski B, Ashburner J & Frackowiak R (2015). Do we need to revise the tripartite subdivision hypothesis of the human subthalamic nucleus (STN)? Response to Alkemade and Forstmann. Neuroimage, 110, 1–2.

51 Lee HK, Braynen W, Keshav K & Pavlidis P (2005). ErmineJ: tool for functional analysis of gene expression data sets. BMC Bioinformatics, 6, 269.

52 Leh SE, Ptito A, Chakravarty MM & Strafella AP (2007). Fronto-striatal connections in the human brain: a probabilistic diffusion tractography study. Neurosci. Lett. 419, 113–8.

53 Lehéricy S, Ducros M, Van de Moortele PF, Francois C, Thivard L, Poupon C, Swindale N, Ugurbil K & Kim DS (2004). Diffusion tensor fiber tracking shows distinct corticostriatal circuits in humans. Ann. Neurol. 55, 522–9.

54 Levey AI, Hersch SM, Rye DB, Sunahara RK, Niznik HB, Kitt CA, Price DL, Maggio R, Brann MR & Ciliax BJ (1993). Localization of D1 and D2 dopamine receptors in brain with subtype-specific antibodies. Proc. Natl. Acad. Sci. U. S. A. 90, 8861–5.

55 Liu J, Hu Q & Yu D (2008). A comparative study on rough set based class imbalance learning. Knowledge-Based Syst. 21, 753–763.

56 Martinez D, Slifstein M, Broft A, Mawlawi O, Hwang Dr, Huang Y, Cooper T, Kegeles L, Zarahn E, Abi-Dargham A, Haber SN & Laruelle M (2003). Imaging human mesolimbic dopamine transmission with positron emission tomography. Part II: amphetamine-induced dopamine release in the functional subdivisions of the striatum. J. Cereb. Blood Flow Metab. 23, 285–300.

57 Meilă M (2007). Comparing clusterings-an information based distance. J. Multivar. Anal. 98, 873–895.

58 Miyachi S, Hikosaka O, Miyashita K, K’ar’adi Z & Rand MK (1997). Differential roles of monkey striatum in learning of sequential hand movement. Exp. brain Res. 115, 1–5.

59 Nelson TH, Jung JY, DeLuca TF, Hinebaugh BK, St. Gabriel KC & Wall DP (2012). Autworks: a cross-disease network biology application for Autism and related disorders. BMC Med. Genomics, 5, 56.

60 Oldham MC, Konopka G, Iwamoto K, Langfelder P, Kato T, Horvath S & Geschwind DH (2008). Functional organization of the transcriptome in human brain. Nat. Neurosci. 11, 1271–82.

61 Olsen CM, Huang Y, Goodwin S, Ciobanu DC, Lu L, Sutter TR & Winder DG (2008). Microarray analysis reveals distinctive signaling between the bed nucleus of the stria terminalis, nucleus accumbens, and dorsal striatum. Physiol. Genomics, 32, 283–98.

62 Parent A & Hazrati LN (1993). Anatomical aspects of information processing in primate basal ganglia. Trends Neurosci. 16, 111–6.

63 Parent A (1990). Extrinsic connections of the basal ganglia. Trends Neurosci. 13, 254–258.

64 Parent A & Hazrati LN (1995). Functional anatomy of the basal ganglia. I. The cortico-basal ganglia-thalamocortical loop. Brain Res. Rev. 20, 91–127.

65 Passingham RE, Stephan KE & Kötter R (2002). The anatomical basis of functional localization in the cortex. Nat. Rev. Neurosci. 3, 606–616.

66 Piray P, den Ouden HE, van der Schaaf ME, Toni I & Cools R (2015). Dopaminergic Modulation of the Functional Ventrodorsal Architecture of the Human Striatum. Cereb. Cortex bhv243.

67 Reep RL & Corwin JV (1999). Topographic organization of the striatal and thalamic connections of rat medial agranular cortex. Brain Res. 841, 43–52.

68 Reep R, Corwin J, Hashimoto A & Watson R (1987). Efferent Connections of the Rostral Portion of Medial Agranular Cortex in Rats. Brain Res. Bull. 19, 203–221.

69 Robbe D, Alonso G & Manzoni OJ (2003). Exogenous and endogenous cannabinoids control synaptic transmission in mice nucleus accumbens. Ann. N. Y. Acad. Sci. 1003, 212–25.

70 Rubinov M & Sporns O (2011). Weight-conserving characterization of complex functional brain networks. Neuroimage, 56, 2068–2079.

71 Schlicker A, Domingues FS, Rahnenführer J & Lengauer T (2006). A new measure for functional similarity of gene products based on Gene Ontology. BMC Bioinformatics, 7, 302.

72 Seeman P & Lee T (1975). Antipsychotic Drugs: Direct Correlation between Clinical Potency and Presynaptic Action on Dopamine Neurons, 188, 1217–1219.

73 Shen EH, Overly CC & Jones, AR (2012). The Allen Human Brain Atlas: comprehensive gene expression mapping of the human brain. Trends Neurosci. 35, 711–4.

74 Simpson EH, Winiger V, Biezonski DK, Haq I, Kandel ER & Kellendonk C (2014).Selective Overexpression of Dopamine D3 Receptors in the Striatum Disrupts Motivation but not Cognition. Biol. Psychiatry, 76, 823–831.

75 Sotiropoulos SN, Jbabdi S, Xu J, Andersson JL, Moeller S, Auerbach EJ, Glasser MF, Hernandez M, Sapiro G, Jenkinson M, Feinberg Da, Yacoub E, Lenglet C, Van Essen DC, Ugurbil K, Behrens TEJ & WU-Minn HCP Consortium (2013). Advances in diffusion MRI acquisition and processing in the Human Connectome Project. Neuroimage, 80, 125–43.

76 Stalnaker Ta, Calhoon GG, Ogawa M, Roesch MR & Schoenbaum G (2010). Neural correlates of stimulus response and response-outcome associations in dorsolateral versus dorsomedial striatum. Front. Integr. Neurosci. 4, 12.

77 Supek F, Bošnjak, M, Škunca N & Šmuc T(2011). REVIGO summarizes and visualizes long lists of gene ontology terms. PLoS One, 6, e21800.

78 Thomas C, Ye FQ, Irfanoglu MO, Modi P, Saleem KS, Leopold DA & Pierpaoli C (2014). Anatomical accuracy of brain connections derived from diffusion MRI tractography is inherently limited. Proc. Natl. Acad. Sci. U. S. A. 111, 16574–9.

79 Tomassini V, Jbabdi S, Klein JC, Behrens TEJ, Pozzilli C, Matthews PM, Rushworth MFS & Johansen-berg H (2007). Diffusion-Weighted Imaging Tractography-Based Parcellation of the Human Lateral Premotor Cortex Identifies Dorsal and Ventral Subregions with Anatomical and Functional Specializations, 27, 10259–10269.

80 Tziortzi AC, Haber SN, Searle GE, Tsoumpas C, Long CJ, Shotbolt P, Douaud G, Jbabdi S, Behrens TEJ, Rabiner, EA, Jenkinson M & Gunn RN (2014). Connectivity-based functional analysis of dopamine release in the striatum using diffusion-weighted MRI and positron emission tomography. Cereb. Cortex, 24, 1165–77.

81 van der Stelt M & Di Marzo V (2003). The endocannabinoid system in the basal ganglia and in the mesolimbic reward system: implications for neurological and psychiatric disorders. Eur. J. Pharmacol. 480, 133–150.

82 Van Essen DC, Smith SM, Barch DM, Behrens TEJ, Yacoub E, Ugurbil K & WU-Minn HCP Consortium (2013). The WU-Minn Human Connectome Project: an overview. Neuroimage, 80, 62–79.

83 Wolf L, Goldberg C, Manor N, Sharan R & Ruppin E (2011). Gene expression in the rodent brain is associated with its regional connectivity. PLoS Comput. Biol. 7, e1002040.

